# Multiscale Transcriptomic Architecture of the ALS Motor Cortex

**DOI:** 10.64898/2026.10.24.761595

**Authors:** Aadya Goel, Shaojun Pei, Gil Alterovitz

## Abstract

Amyotrophic lateral sclerosis (ALS) exhibits marked clinical and molecular heterogeneity, but how this heterogeneity is organized within disease-relevant central nervous system regions remains incompletely understood. Because pooled cross-region analyses may confound regional variation with subtype structure, we focused on the motor cortex, a key site of ALS pathology, to resolve its internal transcriptomic organization. Using approximately 600 postmortem motor cortex transcriptomes from the Target ALS resource, we applied gene set variation analysis to generate sample-level pathway activity profiles and performed consensus clustering in pathway space. We identified seven reproducible ALS motor cortex subtypes characterized by distinct inflammatory, damage-response, metabolic, glial, and neuronal/synaptic programs. Donor-held-out projection recovered the seven subtype assignments with a mean balanced accuracy of 0.939, supporting the reproducibility of the pathway-defined subtype structure in unseen donors. Mapping these bulk-defined programs to an independent motor-cortex single-nucleus RNA-seq dataset revealed distinct cellular contexts, while cell-type marker adjustment showed that cellular differences contributed to, but did not fully account for, subtype-associated pathway variation. The seven subtypes also recapitulated major molecular programs reported in previous ALS subtype frameworks while resolving them into finer motor-cortex-specific states. Rather than forming isolated molecular classes, these subtypes occupied a structured pathway-space landscape with both discrete subtype-centered peaks and continuous biological gradients. Principal component analysis and representative pathway overlays defined a major axis from an upper inflammatory/damage region to a lower neuronal/metabolic/remodeling region, with C7 forming a neuronal/synaptic connector within the lower landscape. Co-expression network analysis further supported this multiscale organization, revealing family-level network backbones, subtype-specific modular refinements, and bridge-like rewiring. Metadata-informed analyses showed that this molecular architecture was not primarily organized by disease duration. Instead, the landscape was more clearly associated with clinical and genetic-background variables, including ALS/FTD family history and mutation-related signals. In particular, family-history-positive and C9orf72 repeat expansion-positive samples were more represented in the lower neuronal/metabolic/remodeling region, whereas family-history-negative samples and C9orf72-negative cases were more represented in the upper inflammatory/damage region. These findings support a model in which ALS motor cortex heterogeneity is organized as a multiscale transcriptomic architecture more closely associated with genetic-background variables than with disease duration. This framework provides a basis for subtype-aware biological comparison, future patient-stratification, biomarker, or therapeutic target-prioritization studies.

## 1 Introduction

Amyotrophic lateral sclerosis (ALS) is a progressive neurodegenerative disease defined by degeneration of upper and lower motor neurons. Patients differ in clinical presentation, anatomical pattern of involvement, rate of progression, cognitive and behavioral symptoms, genetic architecture, neuropathology, and molecular profile, indicating that ALS is better understood as a heterogeneous disease spectrum rather than a single homogeneous entity [1, 2, 3, 8, 9]. This heterogeneity poses a major challenge for mechanistic interpretation and therapeutic development. Defining molecularly coherent ALS subgroups is therefore an important step toward more precise disease characterization and mechanism-informed stratification [2, 3, 4, 11, 12].

Traditionally, ALS heterogeneity has been described using clinical, genetic, and neuropathological classifications [1, 2, 8, 9]. However, although these traditional classifications capture important clinical, genetic, and pathological differences, they do not fully explain the molecular diversity of ALS. Recent transcriptomic and multi-omic studies have therefore extended ALS classification toward molecular subtyping. These studies suggest that unsupervised molecular stratification can identify disease states not fully captured by diagnostic labels or conventional clinical subtypes [3, 11, 12].

A central challenge, however, is that ALS molecular heterogeneity is shaped by anatomical context. As a result, subtype structure inferred from pooled cross-region analyses may reflect anatomical composition as much as disease biology, potentially obscuring or misattributing tissue-specific molecular organization [5, 6, 7, 13, 14, 15]. This makes the motor cortex, a core site of upper motor neuron pathology, a particularly relevant tissue in which to examine ALS subtype architecture within a more coherent anatomical setting [8].

A second challenge is methodological. Transcriptomic subtype discovery is complicated by high dimensionality, biological and technical noise, variation in cell-type composition, sample quality, platform effects, and clinical heterogeneity. Moreover, ALS-associated molecular changes rarely occur as isolated single-gene events. Instead, they involve coordinated biological programs such as RNA processing, proteostasis, mitochondrial metabolism, synaptic function, membrane remodeling, glial activation, and neuroinflammatory response [6, 7, 8, 9, 10]. Patient samples may therefore not form sharply separated and biologically independent classes. They may instead occupy a structured molecular space containing subtype poles, continuous gradients, intermediate states, local overlap, and higher-order families of related disease programs [3, 8, 9, 11, 16]. Conventional clustering labels alone may be insufficient to describe this organization [3, 11, 16].

To address these challenges, subtype analysis needs to move beyond cluster identification alone and characterize the structure of subtype relationships. Pathway-level representations can reduce gene-level complexity and improve biological interpretability by summarizing coordinated molecular activity into disease-relevant programs [6, 7, 10, 13]. At the same time, prior ALS studies have shown how geometric and network-based analyses can reveal how subtype states are arranged, whether they are discrete or continuous, and how pathway-defined states connect to gene-level modules and hub-centered subnetworks [10, 16, 17, 18].

In this study, we examined ALS molecular heterogeneity within the motor cortex using transcriptomes from 600 postmortem samples from the Target ALS resource. We identified reproducible molecular subtypes and reconstructed their relationships within a geometry-informed subtype space. Our goal was not only to identify ALS motor cortex subgroups, but also to determine how those subgroups are organized: whether they form isolated categories, partially overlapping states, or a multiscale transcriptomic architecture containing subtype-specific poles, continuous biological structure, and broader families of related molecular programs.

## 2 Methods

Figure 1 summarizes the overall analytical framework, from tissue-specific subtype discovery through cellular-context analysis, comparison with published ALS frameworks, higher-order landscape and gene-network characterization, and metadata assessment.

**Figure 1:**
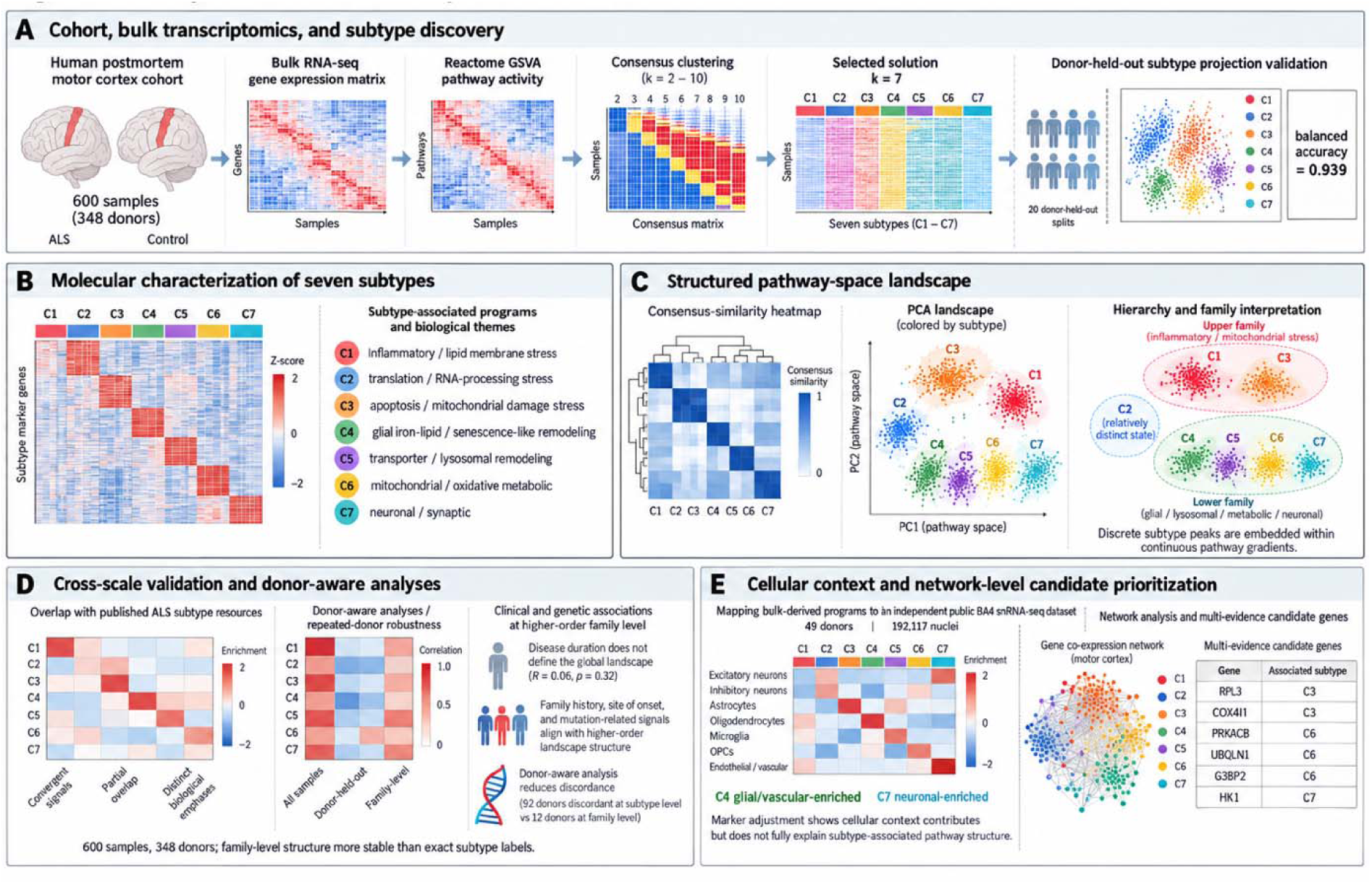
Overall analytical framework for defining and contextualizing ALS motor cortex molecular subtypes. (A) Postmortem motor cortex RNA-seq data were transformed into Reactome pathway-activity profiles using GSVA, and analyzed by consensus clustering across candidate values of k. The selected k = 7 solution defined seven molecular subtypes (C1–C7), whose reproducibility was further evaluated using repeated donor-held-out projection. (B) Subtype-associated pathway and gene-expression patterns defined distinct biological identities. (C) PCA and higher-order relationship analyses showed that these discrete subtype states were embedded within a structured pathway-space landscape containing continuous biological gradients. (D) Cross-scale analyses compared the motor cortex subtypes with previously published ALS molecular subtype frameworks and evaluated clinical, genetic, technical, and donor-level robustness. (E) Bulk-derived molecular programs were mapped to an independent public BA4 single-nucleus RNA-seq dataset to characterize their cellular context. Gene co-expression, marker-gene interaction, and cross-scale network analyses provided complementary support for subtype and family-level biology and prioritized multi-evidence candidate genes for downstream mechanistic investigation. Graphic created by the student researcher using BioRender, 2026.

### 2.1 Motor cortex cohort and gene expression preprocessing

Samples were obtained from the Target ALS Postmortem Tissue Core, a multicenter resource containing molecular and clinicopathologic data from well-characterized postmortem ALS tissue collections [20]. To reduce broad cross-region confounding, the discovery analysis was restricted to 600 postmortem motor cortex samples, spanning BA4, lateral motor cortex, medial motor cortex, unspecified motor cortex, and primary motor cortex. These 600 samples represented 348 unique donors. Repeated donor representation was present with 103 donors contributing one motor-cortex sample, 238 donors contributing two samples, and 7 donors contributing three samples. The selected seven-subtype solution contained C1 = 80, C2 = 50, C3 = 67, C4 = 67, C5 = 119, C6 = 79, and C7 = 138 samples; unique donor representation by subtype was C1 = 64, C2 = 33, C3 = 52, C4 = 53, C5 = 87, C6 = 58, and C7 = 100.

Raw RNA-seq count data were processed using a standardized preprocessing workflow [22–24]. Gene identifiers were harmonized, lowly expressed genes were removed using a CPM-based filter, and counts were normalized for library size and transformed to log2(CPM + 1) values [23–26]. Specifically, genes were retained if they had CPM ≥ 1 in at least 10% of samples. Resulting expression matrix and subtype assignments were examined for residual technical, sample-quality, study/site, and anatomical-region-associated structure after clustering to determine whether these covariates explained the recovered subtype architecture [22, 27]. Because some donors contributed multiple motor-cortex samples, donor identity was retained for downstream sensitivity analyses evaluating donor-level non- independence.

### 2.2 Pathway activity inference and consensus subtype discovery

Normalized gene-expression profiles were transformed into sample-level pathway activity scores using Gene Set Variation Analysis (GSVA) [28], using curated MSigDB collections [29], with Reactome pathways used as the primary functional for its detailed and hierarchical biological annotations [33]. Pathway activity scores were standardized across samples.

Unsupervised consensus clustering was used to evaluate candidate subtype resolutions. For each candidate number of clusters (*K*), samples were repeatedly subsampled, k-means clustering was performed within each subsample, and a consensus matrix was constructed to quantify the stability of pairwise sample co-clustering [30, 31]. Final cluster assignments for each *K* were obtained by hierarchical clustering with average linkage on the consensus-derived distance matrix *D^(k)^ = 1 – C^(k)^*

Candidate solutions were compared using proportion of ambiguous clustering (PAC), consensus cumulative distribution function area, silhouette score, within-cluster distance, cluster-size balance, and representation of motor cortex sampling annotations [30, 32]. Robustness checks compared neighboring clustering solutions and pathway profiles across motor cortex subdivisions. For the donor-held-out recovery, 20% of donors were repeatedly held out, subtype centroids were derived from training samples, and held-out samples were assigned by similarity to these centroids. Recovery was evaluated using top-1 and top-2 agreement against permutation-based null distributions.

Each molecular subtype was biologically characterized using one-versus-rest pathway activity comparisons, subtype-associated genes, and functional enrichment analysis. Pathways were ranked by effect size and statistical evidence, and Reactome-based annotations were used to summarize subtype-defining biological programs [29, 33]. Gene-level contributors to each subtype were then examined in relation to enriched pathways to determine whether subtypes were primarily characterized by neuronal/synaptic programs, immune or inflammatory activity, metabolic alteration, stress response, glial biology, transport remodeling, or other coordinated functional states [34–36]. Functional over- representation analyses were used as post hoc biological annotation rather than as a second clustering procedure. Final subtype labels were assigned after integrating pathway-level enrichment, gene-level contributors, and consistency with known ALS-relevant mechanisms.

### 2.3 Single-nucleus assessment of cell-type-adjusted subtype-defining pathways

Processed single-nucleus RNA-seq count matrices and cell-type annotations were obtained from the publicly available dataset of Pineda et al. [21] (Synapse: syn51105515). Analyses were restricted to BA4 motor cortex samples from 49 donors, including 17 sporadic ALS, 16 C9orf72-associated ALS, and 16 pathologically normal controls, comprising 192,117 nuclei. The seven bulk-derived molecular signatures were mapped to the single-nucleus expression matrix using matched gene identifiers, and signature scores were calculated for individual nuclei and aggregated by annotated cell population and donor. Cellular enrichment of representative signatures was first characterized across excitatory and inhibitory neurons, astrocytes, microglia, oligodendrocytes, oligodendrocyte precursor cells, and vascular cells; within-donor contrasts were evaluated using Wilcoxon signed-rank tests with Benjamini–Hochberg correction. To distinguish shared from subtype-specific cellular signals, overlapping signatures such as C1 and C3 were additionally partitioned into shared and subtype-specific gene components and rescored separately. Robustness of subtype-associated pathway variation was then assessed using nested regression models incorporating marker-expression scores for the major cortical cell populations, together with motor-cortex region and available technical covariates. Subtype-associated variation was quantified using partial *R^2^* before and after inclusion of cell-type marker scores, with robustness further evaluated by equal-donor weighting, 100 one-sample-per-donor iterations, and independent cell-type marker panels.

### 2.4 Comparison with published ALS subtype frameworks

To evaluate how the motor-cortex-derived subtype solution related to previously reported ALS subtype categories, the seven subtypes were compared with a published five-class ALS subtype framework consisting of ALS-Ox, ALS-Glia, ALS-TE, ALS-TD, and ALS-Neu using two complementary signature-scoring strategies. First, published subtype programs were represented in Reactome pathway-activity space using curated pathway signatures. For each published subtype *s*, a pathway-level subtype score was calculated by contrasting the mean standardized activity of pathways assigned to subtype against the mean activity of pathways assigned to the remaining published subtype programs:

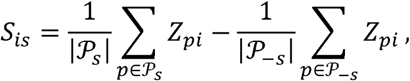

Where *Z_pi_* denotes the standardized GSVA score of pathway *p* in sample *i, P_s_*, denotes the set of pathways assigned to published subtype *s*, and *P_-s_* denotes the set of pathways assigned to the remaining published subtype programs. These scores were used as continuous similarity measures and to identify the closest published subtype program for each sample.

Second, an independent gene-level comparison was performed using curated marker-gene signatures for the same published subtype categories. Gene expression values were normalized, log-transformed, and standardized across samples. For each published subtype *s*, the signed gene-level subtype score was calculated as the mean standardized expression of its marker genes:

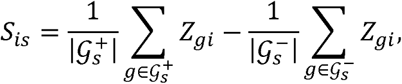

where *Z_gi_* denotes the standardized gene expression value of gene in sample and 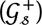 and 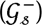 denote the positively and negatively weighted genes in the subtype signature, respectively. For both pathway-level and gene-level scoring, sample-level published subtype scores were summarized within each motor cortex subtype: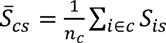, where *n_c_* is the number of samples in cluster *c*.

Differences in published subtype scores across the seven motor cortex clusters were tested using Kruskal–Wallis tests. Cluster-specific enrichment of published subtype programs was evaluated using one-sided Mann–Whitney U tests comparing each cluster against all remaining samples, with Benjamini–Hochberg false-discovery-rate correction.

### 2.5 Subtype landscape and multiscale relationship analysis

Principal component analysis (PCA) was applied to the standardized sample-by-pathway GSVA matrix to visualize the global organization of the seven motor cortex subtypes. The first two principal components were retained as a low-dimensional representation of the pathway-activity landscape, and pathway loadings were examined to identify the biological programs contributing most strongly to each axis.

Five representative Reactome pathways were selected from the major subtype-defining programs: DPAn-6 specialized pro-resolving mediator biosynthesis, BAX activation, AMPA receptor activation, mitochondrial beta oxidation, and organic anion transport by SLC22 transporters. Sample-level activity for each pathway was projected onto PCA space, and its association with PC1 and PC2 was evaluated using Pearson and Spearman correlations. Representative pathway activity was also compared across the seven subtypes using Kruskal–Wallis tests followed by pairwise Mann–Whitney U tests, with false-discovery-rate correction.

To distinguish subtype-centered pathway enrichment from broader continuous landscape variation, pathway activity was modeled using subtype identity, PCA position, or both. The explanatory contributions of subtype membership and PCA position were compared using model ℝ^2^, allowing pathways to be interpreted as primarily subtype-restricted, primarily continuous, or influenced by both forms of organization.

Higher-order subtype relationships were assessed by average-linkage hierarchical clustering using Euclidean distances based on either subtype PCA centroids or mean Reactome pathway profiles. Concordant relationships across these two representations were used to define broader molecular- landscape groups for subsequent network and clinical-association analyses. Donor-held-out prediction was repeated for these higher-order groups.

To quantify continuous structure beyond hard subtype labels, samples were also characterized using soft membership and local neighborhood mixing. Probabilistic membership was estimated from a Gaussian mixture model in PCA space, while a nearest-neighbor graph in the full standardized pathway space was used to quantify the fraction of neighboring samples belonging to each subtype. These measures were used to distinguish subtype-centered samples from samples occupying locally mixed or intermediate positions.

For clinical and genetic analyses, the dominant upper-to-lower molecular-landscape axis was represented as the projection of each sample onto the vector connecting the centroids of the two major landscape regions. If *μ_U_* and *μ_L_* denote the upper and lower centroids, respectively, the unit direction and sample-level landscape score were defined as

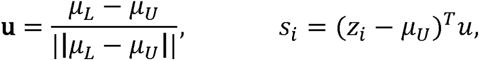

where *z_i_* denotes the PCA coordinates of sample *i*. Higher values therefore represented position toward the lower neuronal/metabolic/remodeling side of the landscape, whereas lower values represented position toward the upper inflammatory/damage side.

### 2.6 Gene-network and pathway-to-module analyses

Gene-level network analyses were used to determine whether the pathway-defined subtype architecture was supported by coordinated gene relationships. Analyses included family-specific co-expression networks, subtype marker-gene interaction networks, and integration of pathway and network evidence.

Within higher-order subtype families, samples close to subtype centers and samples occupying intermediate positions were identified from pathway-space position and subtype-assignment confidence. For each resulting group, lowly expressed genes were removed, the most variable genes were retained, and pairwise Pearson correlations were used to construct sparse weighted co-expression networks from strong positive relationships. Network communities were identified by modularity-based community detection and annotated by Reactome enrichment. Gene importance was summarized primarily by weighted degree. Networks from subtype-core and intermediate groups were compared to identify shared family backbones, subtype-specific organization, and bridge-associated changes in centrality.

Subtype-associated marker genes were identified using one-versus-rest expression comparisons and FDR correction. The highest-ranking markers were used to construct high-confidence STRING interaction networks, from which connected components, communities, and hubs were identified and annotated by Reactome enrichment. Marker, interaction-hub, co-expression-hub, and bridge-centrality evidence were integrated within the relevant higher-order family to prioritize genes supported across multiple analytical layers. Candidate genes were subsequently reviewed using public drug-target and target-annotation resources to distinguish direct target potential from pathway-level or mechanistic relevance.

### 2.7 Metadata-informed analysis across subtype, family, and pathway-program scales

Clinical and genetic-background associations were evaluated at three levels of molecular organization: the seven molecular subtypes, the broader upper C1/C3, C2, and lower C4–C7 landscape regions, and continuous pathway-space features. Available metadata included disease duration, age at symptom onset, age at death, sex, site of motor onset, C9orf72 repeat-expansion status, ALS/FTD family history, reported genomic mutation status, MND with dementia or FTD, and Revised El Escorial classification. Equivalent category labels were harmonized, and unknown, unavailable, or non-informative entries were treated as missing.

For subtype- and region-level analyses, continuous variables were compared using Kruskal–Wallis tests. Categorical variables were evaluated using contingency-table association tests; permutation-based chi-square tests or Fisher’s exact tests were used when expected counts were sparse. Significant global categorical associations were followed by group-versus-rest enrichment analyses using Fisher’s exact tests. P-values were corrected using the Benjamini–Hochberg false-discovery-rate procedure.

Clinical and genetic variables were also evaluated against continuous molecular features derived from the pathway-space analysis, including the upper inflammatory/damage-to-lower neuronal/metabolic/remodeling axis and the representative pathway directions defined in Section 2.5. Continuous clinical variables were tested using Spearman correlation and regression, whereas categorical variables were compared across landscape-feature distributions using Mann–Whitney U or Kruskal–Wallis tests as appropriate.

Disease duration was defined as the reported interval from symptom onset to death. Because all samples were obtained postmortem, these analyses were interpreted as retrospective comparisons of observed disease duration rather than prospective survival prediction. Disease-duration differences were assessed using group comparisons and Cox proportional-hazards models, with additional models adjusted for age at symptom onset and sex.

Because some donors contributed multiple motor-cortex samples, key subtype- and region-level associations were repeated at the donor level. Donors represented by multiple concordant samples were assigned that shared label; otherwise, a majority label was used when available, and donors without a majority assignment were classified as discordant. Sensitivity analyses were performed both excluding discordant donors and retaining them as a separate category.

## 3 Results

### 3.1 Pathway-based consensus clustering identifies seven robust ALS motor cortex subtypes

To define molecular subtype structure within the ALS motor cortex, we performed consensus clustering on the GSVA pathway-activity matrix across candidate resolutions from *k = 2 to 10*. Candidate solutions were evaluated using a prespecified balance of resampling stability, subtype resolution, cluster-size adequacy, and biological interpretability rather than by maximizing a single metric. Among the tested solutions, k = 7 provided the strongest overall balance. Lower-k solutions favored by silhouette compressed distinct pathway programs into broader groups, whereas higher-k solutions showed continued metric improvement in distance-based summaries but introduced smaller and less stable partitions. Thus, k = 7 was selected as the most stable and biologically interpretable solution without evident over-splitting (Figure 2A). The selected *k = 7* solution showed a block-like but nonuniform consensus structure, consistent with the distinct pathway-level profiles observed across the same seven clusters in cluster-versus-rest GSVA comparisons (Figure 2B-2C).

**Figure 2:**
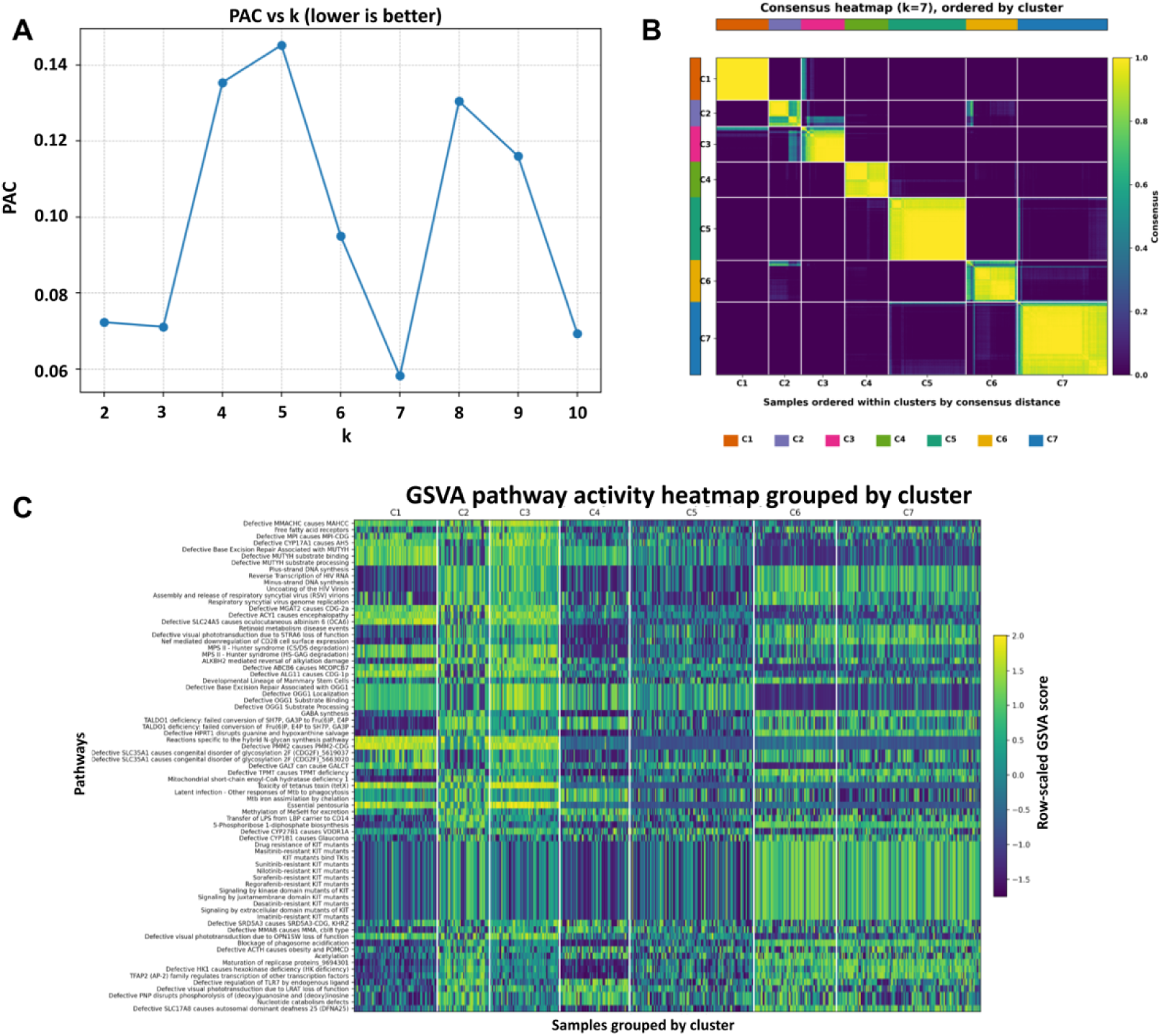
Subtype identification. (A): PAC evaluation of consensus-clustering solutions across *k* = 2 to *k* = 10 for the motor cortex pathway activity matrix. (B) Consensus matrix for the selected *k* = 7 motor cortex solution. (C) GSVA pathway-activity heatmap summarizing the biological identities of the seven motor cortex ALS subtypes. Graphic created by the student researcher using Python, 2026.

The seven subtypes represented distinct but partially related biological programs. C1 was characterized by inflammatory lipid and membrane-remodeling biology, including specialized pro-resolving mediator biosynthesis, membrane-associated lipid processing, inflammasome activity, and stress-related pathways. C2 was dominated by translational and RNA-processing biology, including peptide chain elongation, translation termination, nonsense-mediated decay, and ribosome-rescue programs. C3 showed a damage-response profile centered on redox imbalance, BAX-associated apoptotic activation, metallothionein-related programs, DNA repair-associated defects, and inflammatory stress. C4 was associated with iron handling, lipid transport, and chronic remodeling- like biology, including aceruloplasminemia- and hemochromatosis-related entries and transporter-associated pathways. C5 was enriched for transporter, lysosomal, and metabolic-remodeling programs, including SLC22 organic anion transport, glycosaminoglycan degradation, carnitine transport, and related catabolic processes. C6 represented the clearest mitochondrial and oxidative-metabolism state, with pathways related to mitochondrial beta oxidation, propionyl-CoA catabolism, and broader metabolic dysfunction. C7 captured a neuronal and synaptic signaling state, with enrichment of AMPA receptor activation, GABA synthesis, MECP2-regulated GABA-related transcription, and related neuronal metabolic processes (Figure 2C).

Additional confounding analyses supported the robustness of the selected (k=7) subtype classification. Although anatomical region and technical/sample-quality covariates were associated with cluster membership, neither regional effects nor technical adjustment eliminated the ALS-relevant subtype structure, indicating that the recovered classification was not driven solely by anatomical or technical confounding.

Donor-held-out projection provided an additional test of subtype reproducibility. Across 20 repeated splits, exact C1–C7 assignment achieved a mean accuracy of 0.929, balanced accuracy of 0.939, and macro-F1 of 0.927. The correct subtype was contained within the two highest-ranked assignments for 98.5% of held-out samples. Performance substantially exceeded the corresponding permuted-label null distributions, indicating that the pathway-defined subtype structure was recoverable in donors excluded from model fitting.

### 3.2 Cellular context contributes to but does not fully account for motor cortex subtype structure

Mapping the seven bulk-derived programs to an independent single-nucleus motor-cortex dataset revealed distinct but overlapping cellular distributions (Figure 3). C1 and C2 showed greater representation in endothelial and microglial populations, C3 was enriched in excitatory neurons, and C4 showed prominent vascular, microglial, and astrocytic localization. C5 and C6 were more strongly represented across excitatory neuronal populations, whereas C7 showed the clearest neuronal localization. Donor-level comparisons confirmed the opposing cellular patterns of C4 and C7: C7 scores were consistently higher in excitatory neurons than in glial or vascular populations, whereas C4 scores were higher in glial and vascular cells than in excitatory neurons (FDR ≤ 1.42 × 10^-14^). Analysis of shared and subtype-specific C1/C3 genes further showed that gene overlap contributed to their apparent cellular distributions, emphasizing that the bulk-derived programs should not be interpreted as discrete nucleus-level subtype identities.

**Figure 3:**
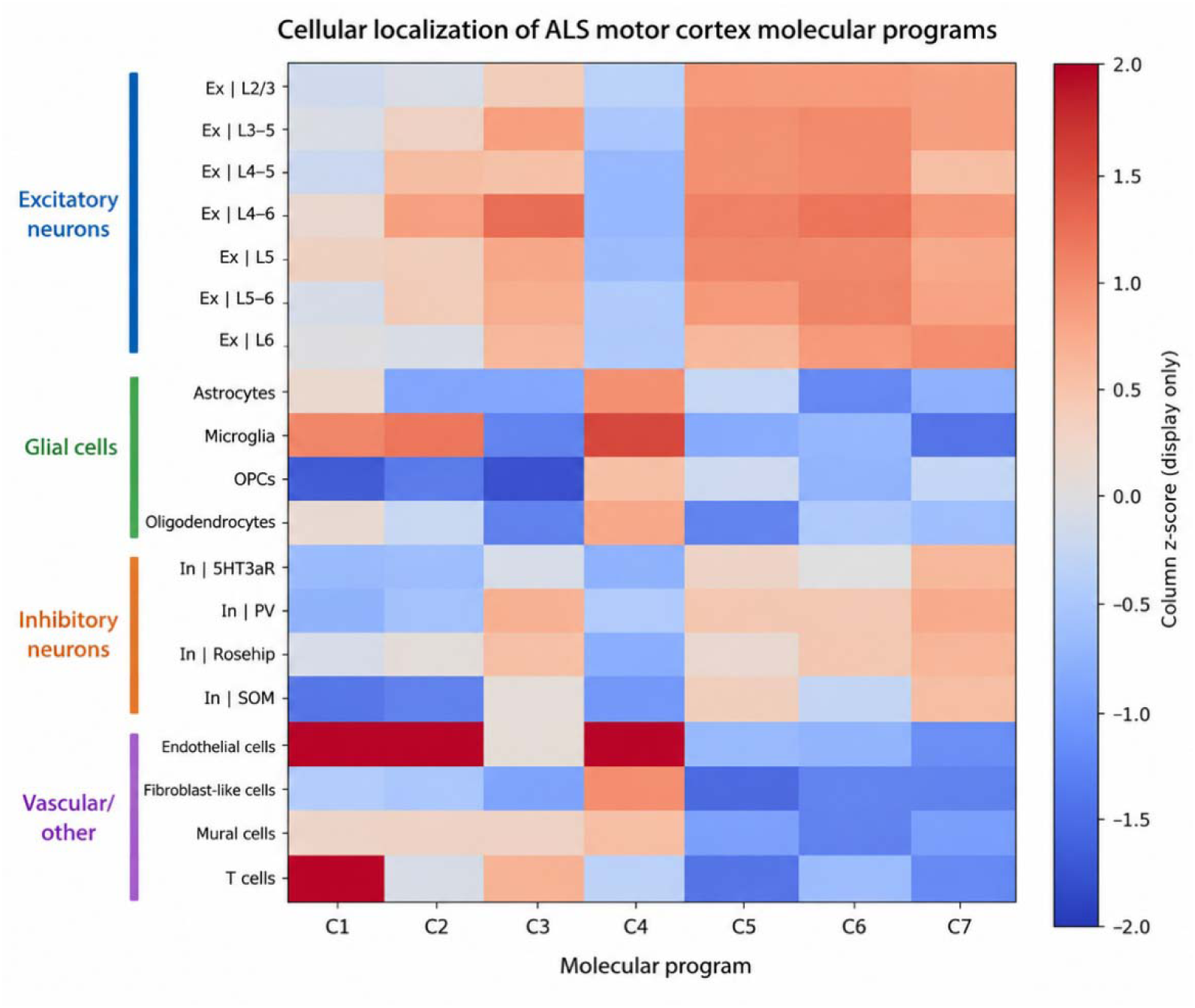
Single-nucleus localization of bulk-derived molecular programs. Relative program scores for the seven bulk-derived molecular programs across annotated cell populations in the BA4 single-nucleus dataset. Graphic created by the student researcher using Python, 2026.

We next asked whether these cellular differences accounted for the subtype-defining pathway structure observed in bulk tissue. Adjustment for major cell-type marker scores substantially attenuated the broad C4 glial/iron-lipid program (median partial R^2^, 0.181 to 0.019) and C7 neuronal/synaptic program (0.110 to 0.003), indicating a strong contribution from cellular context. In contrast, several subtype-associated programs retained appreciable variation after adjustment, including C2 RNA/translation, C3 damage/apoptosis, C5 transport/remodeling, and C6 mitochondrial/metabolic programs. Specific defining pathways also remained prominent, including DPAn-6 specialized pro-resolving mediator biosynthesis in C1 (0.226 to 0.129) and AMPA receptor activation in C7 (0.228 to 0.128). These patterns were consistent across donor-aware and independent-marker-panel sensitivity analyses.

These results indicate that the bulk motor-cortex subtypes capture both differences in cellular context and molecular variation that is not fully explained by broad cell-type-associated expression.

### 3.3 Motor cortex subtypes recapitulate and refine published ALS subtype frameworks

We compared the seven clusters with the five published ALS subtype programs described by Jammal et al.: ALS-Glia, ALS-Neu, ALS-Ox, ALS-TD, and ALS-TE [37]. Two complementary comparisons were performed: a pathway-level comparison based on subtype-associated pathway scores and a gene-signature comparison based on published subtype marker genes (Figure 4). The gene-signature analysis showed strong marker coverage in the motor cortex expression matrix, with complete coverage for ALS-Ox, ALS-Glia, ALS-TD, and ALS-Neu signatures and 37 of 40 ALS-TE genes matched.

**Figure 4:**
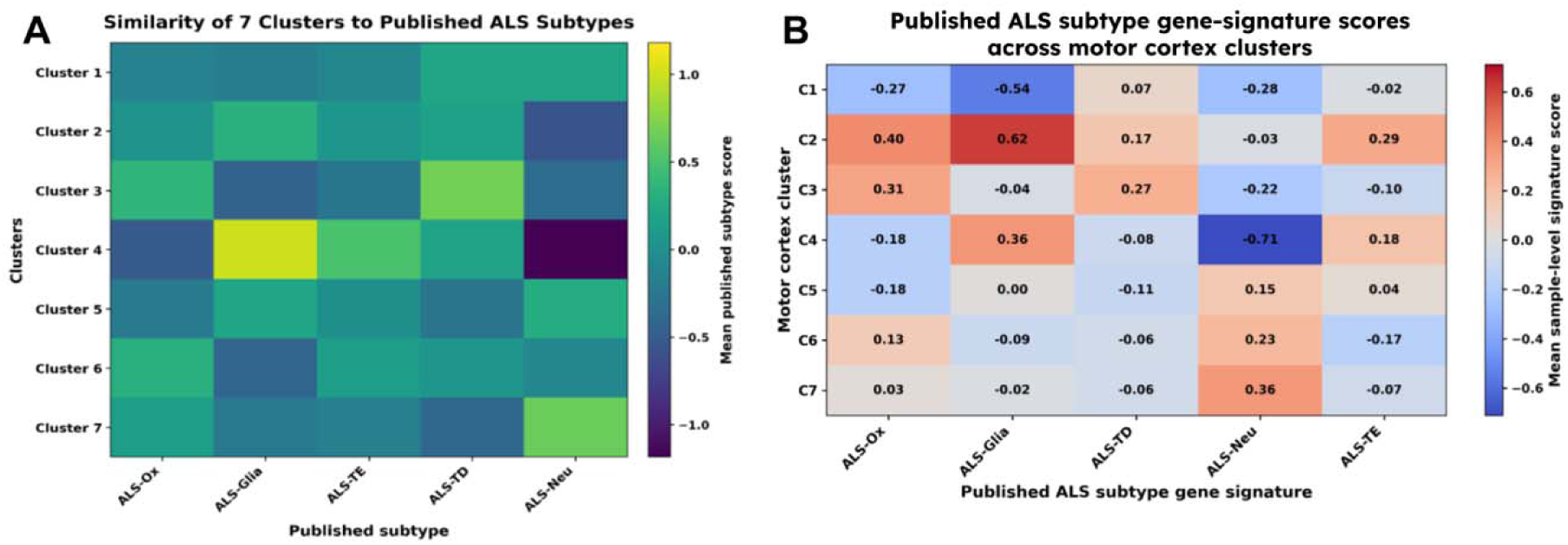
Subtype comparison with published ALS programs. (A) Similarity of seven discovered motor cortex subtypes to published ALS subtype categories in pathway space. (B) Mean published-subtype gene-signature scores across C1-C7. Graphic created by the student researcher using Python, 2026.

The motor cortex subtypes recovered major published programs without mapping one-to-one onto the five-class framework. C7 showed the clearest neuronal (ALS-Neu) alignment. C4 combined glial (ALS-Glia) and TE-associated (ALS-TE) features with reduced neuronal signal (ALS-Neu), whereas C2 combined glial (ALS-Glia), oxidative (ALS-Ox), and TE/interferon-related features (ALS-TE). C3 overlapped oxidative (ALS-Ox) and transcriptional-disruption programs (ALS-TD), consistent with its damage, mitochondrial, and RNA-processing biology. C6 showed weaker oxidative or neuronal-like similarity depending on the scoring layer. C1 and C5 had mixed correspondence. Thus, the motor cortex solution resolved broader published programs into finer, overlapping tissue-specific states.

### 3.4 Higher-order organization of ALS motor cortex subtypes across a continuous pathway-space landscape is supported by gene-network structure

To examine the relationships among the seven motor cortex molecular subtypes, we projected the GSVA pathway-activity matrix into principal-component space. The seven subtypes occupied a structured configuration rather than forming uniformly isolated clusters: C1 and C3 localized to an upper region, C4–C7 occupied a broader lower region, and C2 remained relatively distinct along one side of the landscape (Figure 5A).

**Figure 5:**
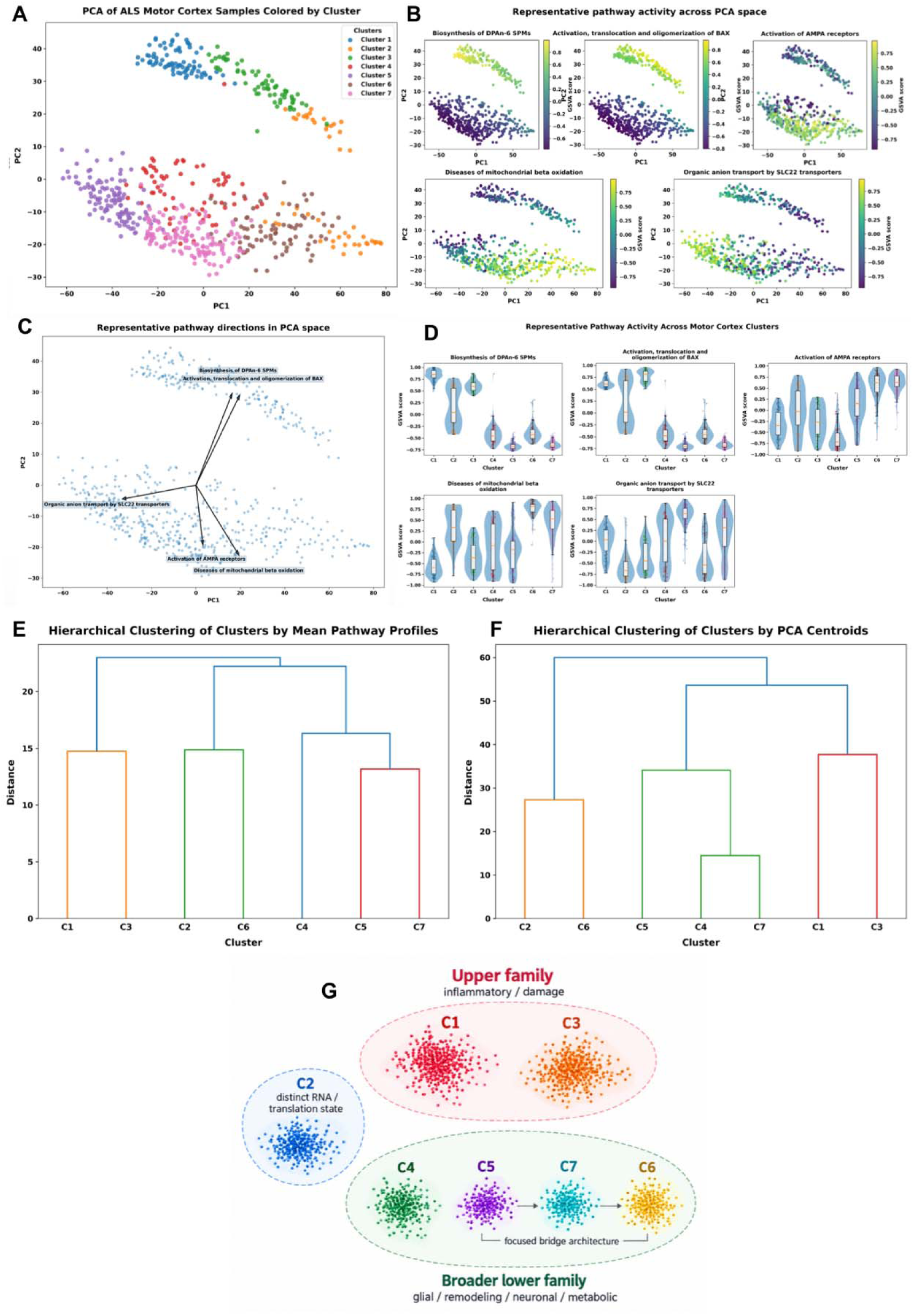
PCA-derived biological landscape of ALS motor cortex subtypes. (A) PCA of ALS motor cortex samples based on the GSVA pathway-activity matrix. (B) Representative pathway activity projected across PCA space. (C) Representative pathway directions in PCA space. Arrows summarize the dominant orientation of major pathway programs across the PCA-derived landscape. (D) Representative pathway activity across the seven motor cortex ALS subtypes. Hierarchical clustering of the seven motor cortex subtypes using (E) mean pathway profiles and (F) PCA centroids. (G) Schematic summary of the higher-order subtype architecture, highlighting the upper C1/C3 family, the distinct C2 state, and the broader lower C4–C7 family, with C5–C7–C6 forming the focused bridge architecture and C4 retained within the broader lower-family structure. Graphic A-F created by the student researcher using Python, 2026. Graphic G created by student researcher using BioRender, 2026.

Representative pathway programs provided biological interpretation of this geometry (Figure 5B– D). DPAn-6 specialized pro-resolving mediator biosynthesis and BAX activation aligned with the upper inflammatory/damage region, with peak activity in C1 and C3, respectively. In contrast, AMPA receptor activation, mitochondrial beta oxidation, and SLC22 transporter activity mapped across the lower landscape and peaked in C7, C6, and C5, corresponding to neuronal/synaptic, metabolic, and transport-remodeling programs. Despite these subtype-centered maxima, pathway activity extended across neighboring states, indicating that discrete subtype-associated programs were embedded within broader continuous pathway gradients.

We next asked whether this landscape organization defined higher-order subtype families. Hierarchical clustering based on both mean Reactome pathway profiles and PCA centroids consistently supported a close C1–C3 relationship and a broader C4–C7 grouping, whereas C2 remained comparatively distinct (Figure 5E-G). Together, these analyses defined an upper C1/C3 inflammatory/damage family, a lower C4–C7 neuronal/metabolic/remodeling family, and a distinct C2 RNA/translation-stress state. This higher-order organization provided the framework for subsequent family-level network and clinical-association analyses.

We next asked whether this higher-order pathway organization was supported at the gene-network level (Figure 6). Family-specific co-expression networks recapitulated the major upper and lower structures. The C1/C3 family contained a shared inflammatory/damage-associated backbone while retaining distinct endpoint organization: C1 emphasized RNA-regulatory, membrane/vesicle, and adhesion programs, whereas C3 showed stronger mitochondrial-stress, DNA-damage, and vesicle/endosomal organization. An intermediate C1/C3 bridge retained features of both endpoint states, supporting a structured relationship between the two subtype programs. Within the lower landscape, C5, C6, and C7 formed transport/remodeling, metabolic/protein-handling, and neuronal/synaptic poles, respectively. C5–C7 and C6–C7 bridge states were identified, whereas no direct C5–C6 bridge was observed, positioning C7 as a connector between remodeling and metabolic programs within the lower landscape (Figure 6A–B).

**Figure 6:**
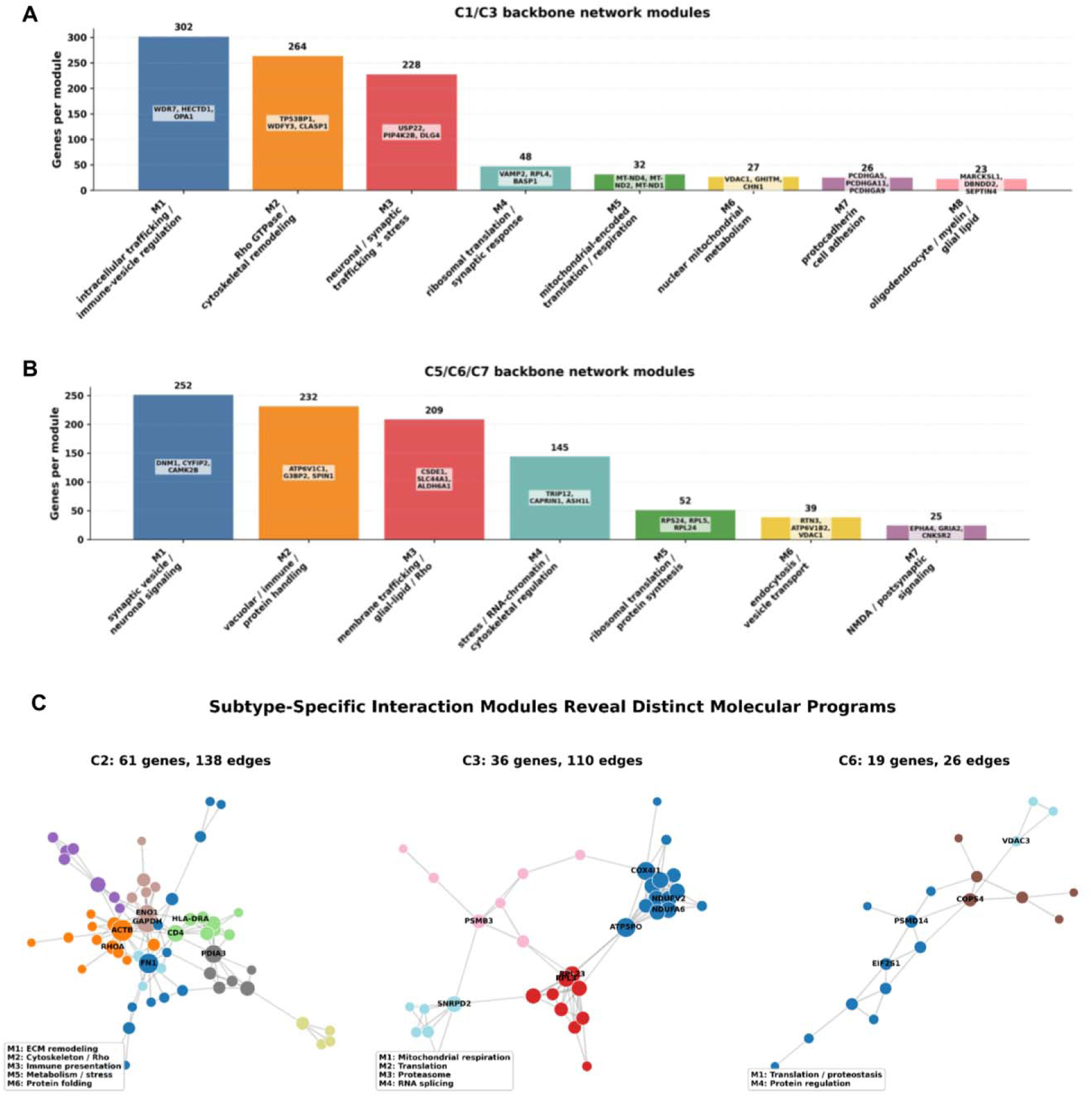
Family-specific co-expression and subtype marker-gene interaction networks. (A-B) Module-size summary for the C1/C3 / C5/C6/C7 backbone network, with modules labeled by dominant biological interpretation and representative hub genes. (C) Major connected components of the C2, C3, and C6 marker-gene interaction networks. The largest connected component is shown for each of the three most organized subtype-level networks. Node size reflects network centrality, node color indicates detected interaction community, and edge width reflects STRING interaction confidence. Selected high-centrality genes are labeled. Graphic created by the student researcher using Python, 2026.

Subtype marker-gene interaction networks provided complementary support for these molecular identities (Figure 6C). C2, C3, and C6 showed the strongest high-confidence interaction structure. C2 contained coordinated immune, extracellular-matrix, antigen-presentation, and protein-stress modules; C3 showed mitochondrial, ribosomal, RNA-processing, and proteostasis organization; and C6 contained mitochondrial-metabolic, translation, proteostasis, and regulatory modules. C7 retained a smaller neuronal/synaptic interaction structure, whereas C1 and C5 were more strongly represented at the pathway and co-expression levels than within curated interaction networks. C2 was therefore retained as a comparatively distinct state rather than incorporated into the family-level bridge analyses. C4 remained part of the broader lower C4–C7 organization but was not represented in the focused C5/C6/C7 bridge architecture.

Finally, integration of subtype-marker, interaction-hub, co-expression-hub, and bridge-centrality evidence identified six genes supported across multiple analytical layers: PRKACB, RPL3, COX4I1, UBQLN1, G3BP2, and HK1. RPL3 and COX4I1 linked the C3 state to translational and mitochondrial programs, PRKACB provided a kinase-associated signal within the C6/lower-family network, and UBQLN1, G3BP2, and HK1 connected subtype-associated expression with proteostasis, stress/RNA regulation, metabolic organization, or bridge-associated network structure. These genes were treated as multi-evidence mechanistic or target-prioritization candidates rather than validated therapeutic targets.

### 3.5 The ALS motor-cortex subtype landscape reflects genetic-background structure rather than disease duration

To determine whether the molecular subtype architecture reflected clinical or genetic-background variation, we evaluated metadata associations across individual subtypes, broader landscape regions, and continuous pathway-space features. Disease duration did not define the global subtype structure. Although median duration varied numerically across the upper C1/C3, C2, and lower C4–C7 regions, these differences were not significant after correction, and continuous landscape features showed no robust association with disease duration (Figure 7A). Thus, the motor-cortex subtype architecture does not appear to represent a simple disease-duration or survival-stage axis.

**Figure 7:**
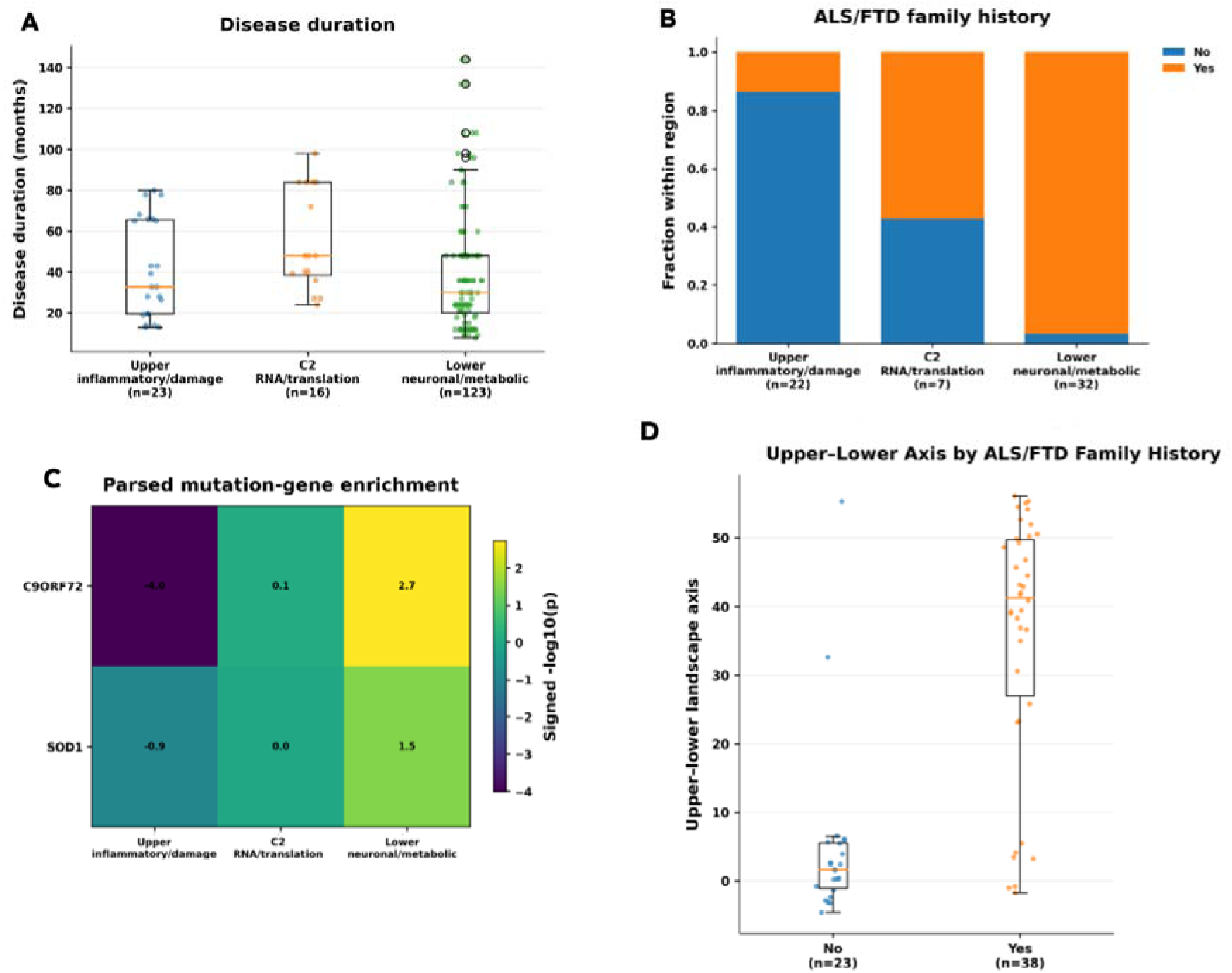
Clinical/genetic-background variation across broader ALS motor-cortex landscape regions. (A) Disease-duration distributions across landscape regions. (B) ALS/FTD family history across landscape regions. (C) Parsed mutation-gene enrichment across landscape regions. (D) Upper-lower landscape- axis scores by ALS/FTD family-history status, showing that family-history-positive samples shift toward the lower neuronal/metabolic/remodeling side of the landscape. Graphic created by the student researcher using Python, 2026.

In contrast, ALS/FTD family history showed a clearer relationship with higher-order molecular organization. Family-history-negative samples were enriched in the upper inflammatory/damage region, whereas family-history-positive samples were more frequently represented in the lower neuronal/metabolic/remodeling region (Figure 7B). Family history also showed the strongest association with the continuous upper-to-lower landscape axis, with family-history-positive samples shifted toward the lower side of the molecular landscape (Figure 7D). This relationship remained after excluding C9orf72-positive samples, indicating that it was not explained solely by C9orf72 repeat-expansion status.

Mutation-related signals showed a similar tendency toward higher-order organization. C9orf72- positive samples were more frequently represented in the lower landscape, while SOD1 showed a weaker lower-region signal (Figure 7C). At the individual-subtype level, C6 showed the clearest SOD1-associated enrichment. Donor-aware analyses supported the broader family-level structure: among 348 donors, discordance decreased from 92 at the seven-subtype level to 12 at the higher-order family level, and ALS/FTD family history remained associated with consensus family assignment. Together, these findings indicate that genetic-background features align more consistently with broader and continuous molecular organization than with individual subtype labels.

## 4 Discussion

The seven subtypes captured major ALS-relevant biological programs, including inflammatory lipid and membrane remodeling, RNA-processing and translational stress, apoptosis and damage response, glial/iron-lipid remodeling, transporter and lysosomal remodeling, mitochondrial and oxidative metabolism, and neuronal/synaptic signaling. Importantly, these pathway-defined differences also reflected distinct cellular contexts, indicating the motor cortex subtype structure is shaped jointly by cellular context and pathway-level molecular variation. Comparison with published ALS subtype frameworks showed that the motor cortex solution recovered major neuronal, glial, oxidative, transcriptional/RNA-processing, and TE/interferon-associated programs, but did not map one-to-one onto previous classifications [2,3,11,37]. This suggests that broad ALS molecular categories can be refined into more specific tissue-defined states when analyzed within a coherent anatomical context. Thus, our results should be viewed not as replacing prior ALS subtype frameworks, but as extending them by resolving motor-cortex-specific structure.

A major conceptual advance of this work is the use of subtype geometry to move beyond hard clustering. The seven motor cortex subtypes contained recognizable biological peaks, but these peaks were embedded in broader gradients of pathway activity. For example, inflammatory lipid and damage-response programs were concentrated in the upper C1/C3 region, whereas neuronal, metabolic, and remodeling programs were distributed across the lower C5/C6/C7 region. The stronger donor-held-out recovery observed at the broader landscape-region level further supports this hierarchical organization, suggesting that individual subtype states are embedded within more stable higher-order molecular neighborhoods. Thus, ALS motor cortex subtype identity appears to contain both categorical and continuous components. This has practical implications for future ALS stratification. Future subtype assignment models may therefore need to move beyond hard labels and incorporate probabilistic membership or continuous landscape position. Such a model is consistent with emerging evidence that ALS molecular subtypes can reflect combinations of cellular and pathological features rather than mutually exclusive disease classes [37,38].

The family-specific networks connected pathway-level organization to gene-level mechanisms. The upper C1/C3 family shared inflammatory and damage-associated biology while retaining distinct endpoint programs, whereas within the lower family, transport/remodeling and metabolic states connected through the neuronal/synaptic C7 neighborhood. Cross-scale integration further identified six multi-evidence candidate genes—PRKACB, RPL3, COX4I1, UBQLN1, G3BP2, and HK1— supported across subtype-marker, interaction-network, co-expression, and bridge-centrality analyses. These candidates linked the broader subtype architecture to specific processes involving translation and mitochondrial respiration, kinase signaling, proteostasis and stress/RNA regulation, and metabolic organization and were prioritized for future investigation.

Unlike previous ALS subtyping studies that reported survival or duration associations [3], this motor cortex landscape showed limited global association with disease duration and clearer associations with family history and genetic background. This suggests a different dimension of ALS heterogeneity, rather than an absence of clinical relevance: region-specific molecular states may reflect genetic-background structure more strongly than a simple survival-stage axis.

Several limitations should be considered. First, Postmortem transcriptomes capture late-stage disease biology at a single time point, so bridge-like organization cannot establish temporal transitions between subtype states. Second, cell-type marker scores measure associated expression rather than direct cellular abundance. Thus, attenuation of selected programs cannot distinguish differences in cell composition from altered transcriptional states within those cell populations. Independent single-nucleus or spatial cohorts are needed to resolve these mechanisms. Third, anatomical and technical covariates were associated with cluster membership, although sensitivity analyses indicated that neither fully explained subtype structure. Future work should validate these subtypes in independent cohorts and develop predictive models that preserve both subtype identity and continuous landscape position.

## 5 Conclusion

In conclusion, this study defines ALS motor cortex heterogeneity as a multiscale transcriptomic architecture rather than a single disease state or a set of isolated molecular clusters. We identified seven reproducible pathway-based subtypes organized within a structured landscape, where discrete subtype identities coexist with broader biological gradients, subtype families, and bridge-like relationships. Gene-network analyses further supported this organization by revealing shared family backbones, subtype-specific refinements, and multi-evidence candidate genes. The architecture refines published ALS frameworks and aligns more clearly with genetic-background features than global disease duration. These findings provide a foundation for subtype-aware biological interpretation and for future biomarker or target-prioritization studies informed by these associations.

## Ethics, Consent to Participate, and Consent to Publish declarations

Not applicable.

## Authors’ contributions

AG and SP conceived and designed the study. AG performed the data analysis and drafted the manuscript. AG and SP contributed to methodology development and interpretation of results. GA provided critical resources/data and supervised the project. All authors read, revised, and approved the final manuscript.

## Acknowledgements

I would like to thank the Target ALS Postmortem Tissue and Genomics Core for providing me with the data used in this research. I would also like to thank my mentors, Dr. Alterovitz and Dr. Pei, for all their help and support throughout this process, alongside the MIT PRIMES programs for providing me with this opportunity.

